# Using *spIsoNet* to address the preferred-orientation problem in cryoEM reconstructions

**DOI:** 10.64898/2026.06.29.735357

**Authors:** Hongcheng Fan, Yun-Tao Liu, Z. Hong Zhou

**Affiliations:** Department of Microbiology, Immunology, and Molecular Genetics, University of California, Los Angeles, CA, USA; California NanoSystems Institute, University of California, Los Angeles, CA, USA

## Abstract

Cryogenic electron microscopy (cryoEM) is now routinely used for high-resolution structure determination of biological macromolecules. However, many biological specimens exhibit varying degrees of preferred orientation on cryoEM grids, resulting in uneven sampling of three-dimensional Fourier space. This orientation bias produces anisotropic reconstruction artifacts and, in severe cases, can exacerbate particle misalignment during iterative refinement, thereby limiting the success rate of near-atomic resolution cryoEM structure determination. This protocol provides a practical guide for applying *spIsoNet*, a self-supervised deep-learning method, to mitigate preferred-orientation issues in cryoEM reconstructions. We describe two complementary workflows: (1) map Anisotropy Correction to correct anisotropic artifacts of cryoEM maps and (2) particle Misalignment Correction, which integrates *spIsoNet* with RELION external reconstruction to improve particle-pose estimation. We demonstrate these workflows using two influenza hemagglutinin (HA) trimer datasets representing moderate and severe degrees of preferred-orientation bias. The protocol includes installation instructions, parameter-selection guidance, quality-control checkpoints and troubleshooting advice, and can typically be completed in ∼7 hours on a workstation equipped with four NVIDIA A100 GPUs. Together, these workflows provide step-by-step guidance for using the open-source *spIsoNet* software to mitigate the preferred-orientation problem directly from experimental data.

## Introduction

### The preferred-orientation problem in cryoEM

Single-particle cryogenic electron microscopy (cryoEM) is widely used in structural biology to resolve high-resolution structures of biological macromolecules in near-native states. Uniformly distributed particle orientations in vitrified ice are critical for complete Fourier-space sampling and for achieving isotropic resolution in single-particle cryoEM reconstructions^1, 2^. However, during cryoEM sample preparation, specimens often adsorb preferentially to the air-water interface (AWI) or the support-film interface, leading to preferred orientations. Such specimen adsorption can bias particle orientations (i.e., “preferred orientation”) and may also promote partial dissociation or even denaturation of biological macromolecules^3^.

Preferred orientation causes missing or under-sampled regions in Fourier space, leading to direction-dependent resolution and anisotropic reconstruction artifacts. In real-space cryoEM maps, this anisotropy can be apparent as density that appears well defined from one viewing direction but streaky, elongated or poorly resolved from orthogonal directions. Typical manifestations include overall density elongation, distorted secondary-structure features, and discontinuous side-chain or nucleobase densities^4^. When an anisotropic map is used as the reference during iterative particle refinement, pose-estimation errors may accumulate, causing the refinement to converge to a suboptimal reconstruction or even fail^5, 6^. Therefore, a general, specimen-independent method remains highly desirable for routine high-resolution cryoEM structure determination and downstream structure-guided studies.

### Methods available to assess, alleviate or mitigate the problem

Several computational tools are available for assessing preferred orientation and its consequences in cryoEM reconstructions. Map-based metrics evaluate anisotropy directly from the reconstructed map. For example, the three-dimensional Fourier shell correlation (3DFSC) metric estimates directional variations in resolution and reports map anisotropy using 3DFSC sphericity^5^; more recently, Fourier shell occupancy (FSO) and the Bingham test (BT) have been introduced as complementary descriptors of resolution anisotropy in cryoEM maps^7^. The FSO curve is evaluated across increasing spatial-frequency shells, with values close to 1 indicating near-uniform isotropy and lower values indicating increasing anisotropy. BT complements FSO by testing whether the reliable directions are uniformly distributed on the sphere, helping distinguish directional anisotropy from a more uniform loss of high-resolution signal^7^.

Orientation-distribution-based methods, such as cryoEF, instead assess preferred orientation from assigned particle Euler angles. CryoEF computes an orientation-distribution efficiency metric, Eod, which characterizes how well a given angular distribution can provide uniform Fourier-space coverage and isotropic resolution. Lower Eod values indicate that non-uniform angular sampling is more likely to produce direction-dependent resolution loss and anisotropic features in the reconstruction^8^. These tools are useful for diagnosis, but additional correction strategies are needed when the goal is to improve reconstructions affected by the underlying orientation bias.

Toward this end, earlier efforts focused on alleviating the preferred orientation problem through experimental strategies. They include chemical modification of specimens^9^, the addition of surfactants to reduce particle adsorption at the air–water interface (AWI)^10–13^; and physical approaches such as the use of cryoEM grids coated with continuous support films^3, 14–18^, time-resolved vitrification devices^19–21^, data collection from thicker ice regions, and tilted-stage imaging^5, 22, 23^. These approaches can be effective in individual cases, but they are often time-consuming and require additional trial-and-error sample screening for optimization. They may be accompanied by trade-offs, including increased background signal, altered particle behavior at newly introduced support-water interfaces, exacerbated beam-induced motion, or larger defocus gradients in tilted datasets. Computational methods therefore provide a complementary route, particularly for datasets that have already been collected or for samples for which experimental re-optimization is difficult.

Recent efforts have led to the development of computational methods that mitigate reconstruction artifacts associated with preferred orientation. Our group developed *spIsoNet*^4^ that provides an end-to-end, self-supervised deep learning method for processing preferred-orientation-limited single particle cryoEM and cryogenic electron tomography (cryoET) subtomogram averaging (STA) datasets. The method is implemented through two workflows to overcome the preferred orientation problem: (1) in the map Anisotropy Correction workflow, users start from half-maps, a solvent mask and an anisotropy descriptor, such as a 3DFSC volume, to generate corrected half-maps for downstream map post-processing and model building; and (2) in the particle Misalignment Correction workflow, *spIsoNet* is integrated with RELION external reconstruction so that *spIsoNet-*corrected half-maps can guide particle pose estimation during iterative refinement. Similar to the idea of the Misalignment Correction workflow of *spIsoNet*, *cryoPROS*^24^ tackles the preferred orientation problem by overcoming mis-alignment with evenly sampled templates called “auxiliary particles”. It uses a self-supervised deep generative model to generate such auxiliary particles with more balanced orientation distributions and co-refines them with experimental particles to improve pose estimation in single-particle cryoEM^24^. Along a similar principle of recovering missing information, *AR-Decon*^25^ uses deconvolution of a point spread function that captures the preferred orientation of the sample to improve map anisotropy without using a deep learning framework. Two other computational tools, *DeepEMhancer*^26^ and *EMReady*^27^ are supervised deep-learning map post-processing tools trained on experimental cryoEM maps paired with model-informed target densities, and are subsequently applied to enhance the visual interpretability of cryoEM maps of interest. In addition, the signal-to-noise ratio iterative reconstruction method (SIRM) aims to mitigate preferred-orientation induced map distortions by removing misaligned particles, followed by dual-space-constrained iterative reconstruction guided by the signal-to-noise ratio of Fourier components^28^.

Since the publication of the original *spIsoNet* paper in 2025^4^, the method has attracted widespread interest, as reflected by its growing number of citations. Nonetheless, many users have expressed interest in a more accessible illustration of its practical implementation in a representative use case. Here, we provide a detailed operational protocol for two *spIsoNet* workflows, each covering input preparation, parameter selection, quality-control checkpoints, troubleshooting and downstream validation.

### Applications

The *spIsoNet* package comprises two modules: (1) map Anisotropy Correction and (2) Anisotropy Correction-powered particle Misalignment Correction. These modules can be applied independently or sequentially, depending on whether the primary failure mode is anisotropic map distortion, particle pose estimation errors, or both. When both map anisotropy and pose estimation errors are evident, we recommend applying particle Misalignment Correction first, followed by map Anisotropy Correction to further improve map isotropy and interpretability.

*SpIsoNet* can be applied to both single particle analysis (SPA) in cryoEM and subtomogram averaging (STA) in cryoET. In SPA, *spIsoNet* has been used to improve map isotropy and particle-pose estimation in datasets affected by preferred orientation. A representative example is the previously intractable untilted influenza hemagglutinin (HA) trimer dataset EMPIAR-10096, which yielded a near-atomic resolution reconstruction after *spIsoNet* particle Misalignment Correction. In STA, *spIsoNe*t has also been shown to improve reconstructions when non-uniform particle orientations compromise map quality^4^. Beyond these benchmark datasets, several cryoEM and cryoET studies have used *spIsoNet* to improve map quality and interpretability in preferred-orientation biased datasets^29–33^.

### Limitations

*spIsoNet* does not generate high-resolution information beyond what is supported by the experimental cryoEM data. The Misalignment Correction workflow is also more computationally demanding than standard RELION refinement because it is embedded in an external-reconstruction loop, in which the *spIsoNet* model is retrained at each RELION refinement iteration. For cryoEM datasets with extremely limited angular coverage, such as those dominated by one or a few preferred views with substantial under-sampling in Fourier space, the ability of *spIsoNet* to improve map isotropy may be limited. In such cases, the situation can be mitigated by combining *spIsoNet* with a straight-forward experimental strategy to alleviate the severe preferred-orientation problem, such as using a single tilt angle within the 20-30° range^5^, using cryoEM grids coated with a continuous support film^3^, or adding a mild detergent to the sample^10^.

### Overview of the *spIsoNet* pipeline

This protocol presents practical workflows established through testing across multiple cryoEM datasets (Fig. 1 and Table 1). Procedure 1, map Anisotropy Correction, takes as input (i) two cryoEM half-maps (Fig. 2a-b), (ii) a solvent mask (Fig. 2c-d), and (iii) a descriptor of directional-resolution anisotropy, for which we used a 3DFSC volume (Fig. 2e-f). A U-Net-style model is trained in a self-supervised manner to generate anisotropy-corrected half-maps that can be used for downstream post-processing, visualization, and atomic-model building. Procedure 2, particle Misalignment Correction, incorporates Anisotropy Correction into an iterative RELION auto-refine workflow. After each refinement iteration, the half-maps are corrected using *spIsoNet* and then low-pass filtered to the current gold-standard FSC resolution before being used as reference maps for the next refinement iteration. The final output is a STAR file with updated particle Euler angles, together with a pair of RELION reconstructed half-maps suitable for downstream postprocessing, visualization and model building. These half-maps can be further subjected to an additional round of Procedure 1 for further map Anisotropy Correction (Fig. 1).

**Fig. 1.**
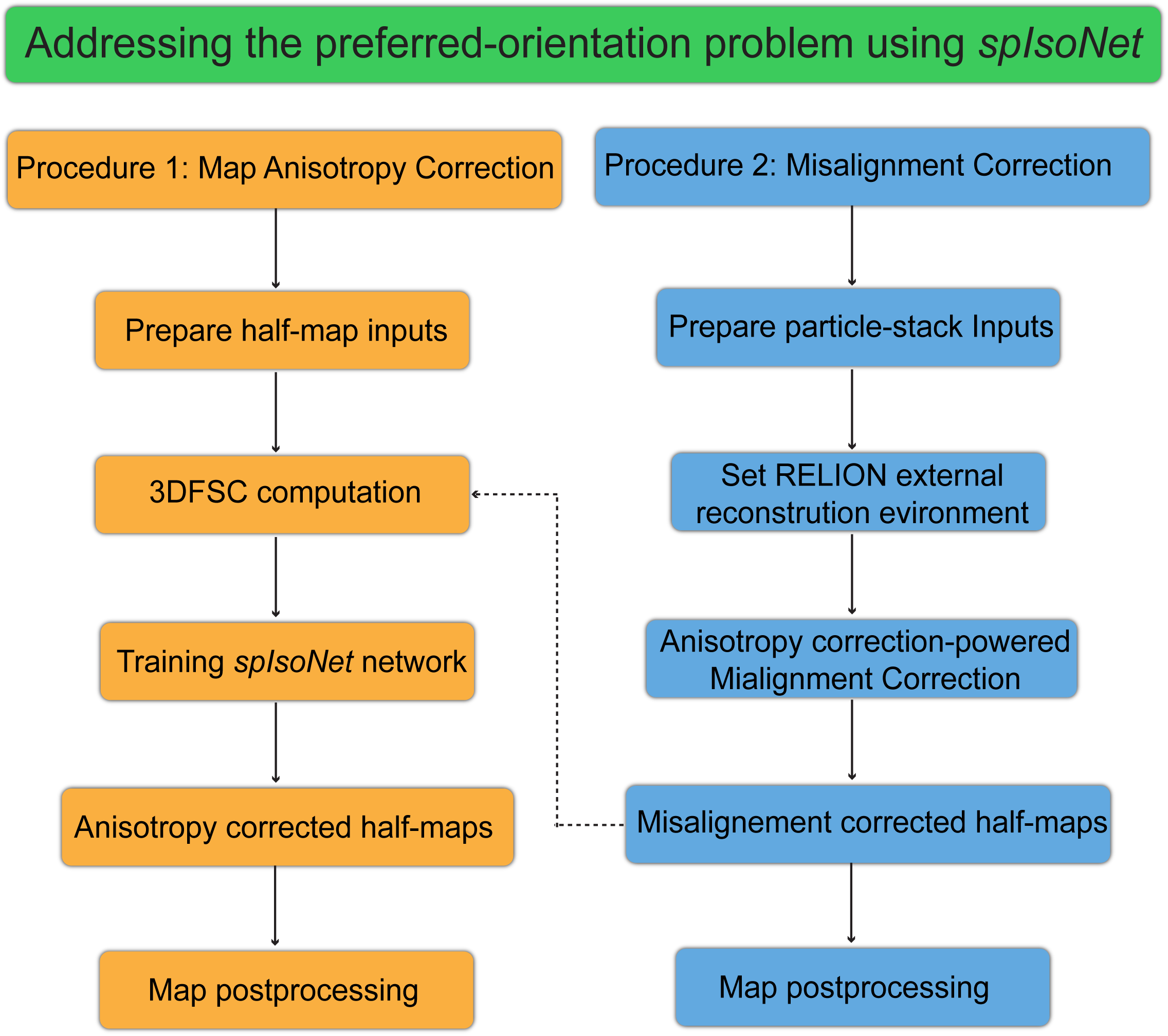
Workflows for cryoEM data processing with *spIsoNet*. Two complementary procedures are shown: Procedure 1, Map Anisotropy Correction (orange), and Procedure 2, Particle Misalignment Correction (blue). In Procedure 1, two independent half-maps are prepared and a 3DFSC volume is computed in *spIsoNet*. *spIsoNet* is then trained in a self-supervised manner using the half-maps and the 3DFSC volume to finally produce anisotropy-corrected half-maps, which are subsequently subjected to standard map post-processing. In Procedure 2, raw particle stacks and associated metadata file are prepared, and an external reconstruction environment is configured in RELION so that *spIsoNet* can be called during iterative refinement. Each refinement iteration uses *spIsoNet*-based Anisotropy Correction to regularize the reference maps, improving particle orientation assignment and finally yielding misalignment-corrected half maps for downstream map post-processing. The optional dashed connection indicates that misalignment-corrected half-maps from Procedure 2 can be used as input for a further round of Anisotropy Correction using Procedure 1. Together, the two procedures mitigate two major consequences of preferred orientation: particle misalignment and map anisotropy.

**Fig. 2.**
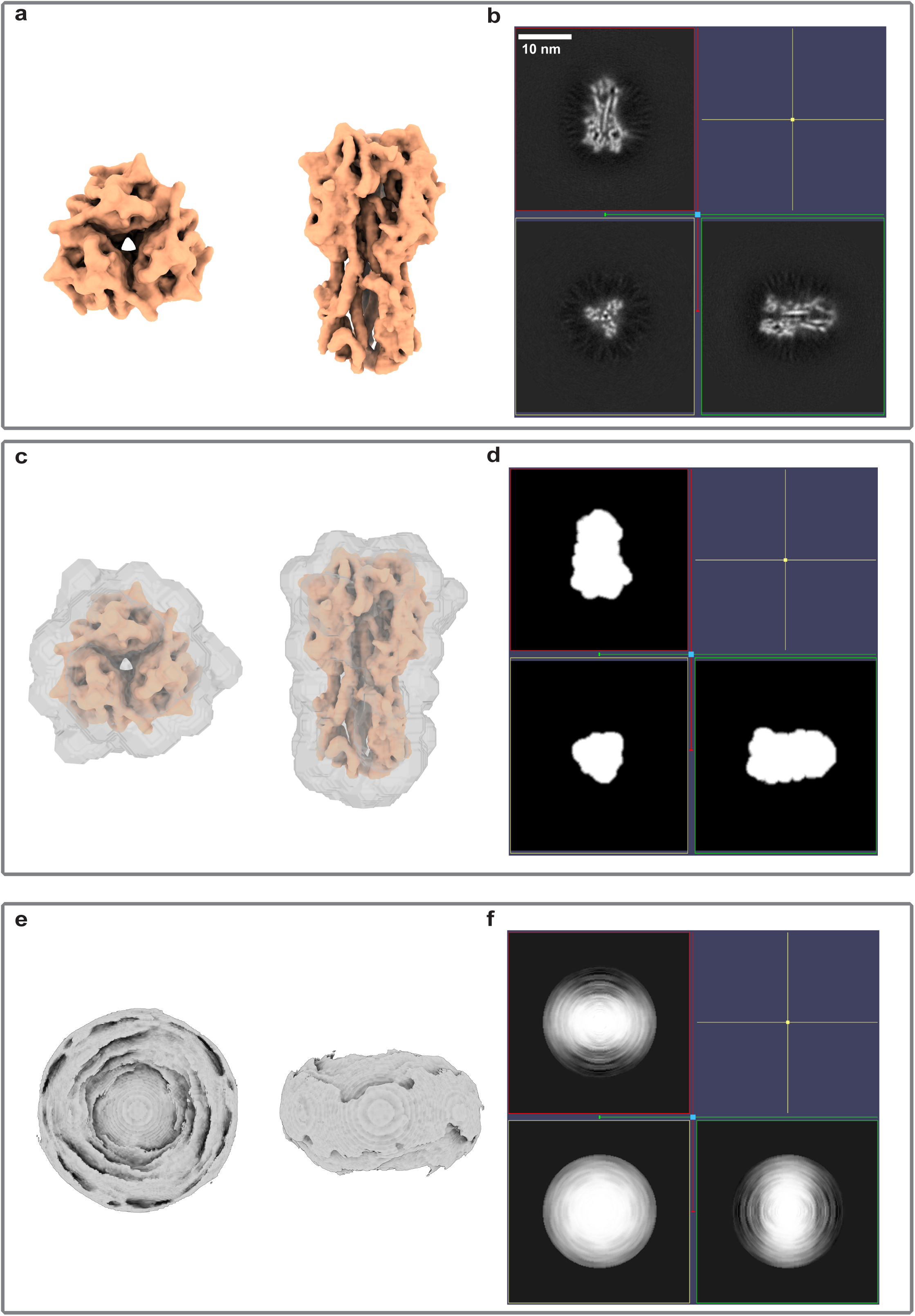
Input files required for *spIsoNet*. **a**, Representative cryoEM half-map. **b**, Orthogonal slice through the cryoEM half-map. **c**, Representative cryoEM half-map overlaid with the solvent mask. **d**, Orthogonal slice through the solvent mask. **e**, Representative 3DFSC volume. **f,** Orthogonal slice through the 3DFSC volume.

**Table 1.** ***SpIsoNet* workflows**

| Workflow | Required inputs | Main tool | Main outputs | Use case |
| --- | --- | --- | --- | --- |
| Map Anisotropy Correction | Two half-maps, a solvent mask and a 3DFSC volume | spisonet.py<br>reconstruct | Anisotropy-corrected half-maps, a trained model, and the loss curve | Maps affected by directional resolution anisotropy |
| Particle Misalignment Correction | A Particle stack, particle metadata file with CTF information, a reference map, and a solvent mask | RELION external reconstruction with *spIsoNet* | Updated particle poses and RELION generated half-maps | Reconstruction artifacts affected by the particle misalignment |

Together, these workflows provide a guided computational strategy for addressing two common consequences of preferred orientation: direction-dependent map anisotropy and particle pose-estimation errors during iterative refinement. We expect that the approach described here will be broadly applicable to the processing of diverse cryoEM datasets.

### Prerequisite for using the protocol

This protocol is intended for cryoEM projects in which preferred orientation impedes structure determination. *spIsoNet* supports multi-GPU acceleration and parallelized processing to improve computational efficiency. Because *spIsoNet* is command-line based, users should be comfortable working in a Unix/Linux environment and should have basic experience with cryoEM data processing, particularly RELION-based refinement for the Misalignment Correction workflow. Users can also refer to the associated *spIsoNet* tutorial, provided in Supplementary Information 1 and available through the *spIsoNet* GitHub repository (https://github.com/IsoNet-cryoET/spIsoNet/tree/main/tutorial), for a more detailed explanation of the algorithms underlying each step, as well as instructions for running the workflows.

## Materials

### Equipment and setup Hardware

The minimum computational requirements for running this protocol include a Linux-based workstation or computing cluster equipped with at least one NVIDIA GPU with at least 12 GB of GPU memory, at least 64 CPU cores, 64 GB of random-access memory (RAM), and 250 GB of available disk space for storing raw data and output files. These requirements are comparable to those of conventional cryoEM reconstruction software.

### Software

*spIsoNet* version 1.0, distributed under the MIT license, is required. We recommend installing *spIsoNet* in an Anaconda or Miniconda environment. Detailed installation instructions are maintained at the *spIsoNet* GitHub repository (https://github.com/IsoNet-cryoET/spIsoNet) and are also provided in Supplementary Information 1.

RELION version 3.1 or 4.0 is required and is available from http://www2.mrc-lmb.cam.ac.uk/relion or https://github.com/3dem/relion.

IMOD version 4.11 or later is required and is available from http://bio3d.colorado.edu/imod/.

UCSF ChimeraX version 1.6.0 or later is required and is available from https://www.cgl.ucsf.edu/chimerax/download.html.

### Datasets

EMD-8731 cryoEM half-maps can be downloaded from https://zenodo.org/records/20783378 or through the EMDB web interface at https://www.ebi.ac.uk/emdb/EMD-8731/

The EMPIAR-10096 subset particle stack can be downloaded from https://zenodo.org/records/20783378. The full particle stack can be downloaded through the EMPIAR web interface at https://www.ebi.ac.uk/empiar/EMPIAR-10096/.

The reference map and associated particle metadata file, in STAR format, used for *spIsoNet* particle Misalignment Correction of EMPIAR-10096 can be downloaded from https://zenodo.org/records/20783378.

All expected results and tutorial data for this protocol can be downloaded from https://zenodo.org/records/20783378.

## Procedure

### CRITICAL

This protocol provides step-by-step procedures for applying *spIsoNet* through two modules: Procedure 1, map Anisotropy Correction using cryoEM half-maps only, and Procedure 2, particle Misalignment Correction by integrating *spIsoNet* into RELION external reconstruction. Map Anisotropy Correction can be used as a standalone workflow, whereas particle Misalignment Correction, requires RELION external reconstruction. We encourage users to first run Procedure 1 as a quick test to evaluate the improvement of their own cryoEM maps after map Anisotropy correction. However, particle alignment may still be affected by map distortion caused by preferred orientation. Procedure 2 is therefore useful for improving particle alignment. These two procedures can be used either independently or in tandem; for example, users may first perform particle Misalignment Correction and then apply map Anisotropy Correction to address two major ramifications of the preferred-orientation problem.

User should run all commands in a terminal shell within the project directory. Users should modify and optimize key parameters according to their own projects. We provide individual commands with specific parameters, discuss the expected results, and describe potential issues and troubleshooting strategies. Novice users are encouraged to follow the steps sequentially, inspect the output from each step, and compare their results with those described here. A more comprehensive tutorial is provided in Supplementary Information 1 and is also available at https://github.com/IsoNet-cryoET/spIsoNet/tree/main/tutorial.

**Procedure 1| Map Anisotropy Correction** ● Timing ∼40 min

**Preparation: arrangement of input files and directories** ● Timing ∼1 min

1 Connect to the workstation on which *spIsoNet* is installed and all required files have been downloaded. We assume that the base directory .`/spIsonet_anisotropy/EMDB-8731/` contains the downloaded EMDB-8731 half-maps (`emd_8731_half_map_1.mrc` and `emd_8731_half_map_2.mrc`), the solvent mask (`emd_8731_msk_1.mrc`), and the target map (`emd_8731.mrc`) with anisotropic resolution that will be improved in this procedure. Most commands in this procedure are run from this base directory. Ensure that the *spIsoNet* conda environment is activated, as this is required for all *spIsoNet* commands. Enter the following commands at the terminal:

`cd /path/to/base/directory`

`conda activate spisonet`

`spisonet.py --help`

The final lines provide a list of available *spIsoNet* commands. To learn more about a specific *spIsoNet* command, simply enter:

`spIsoNet.py [command] --help`

### PAUSE POINT

The two half-maps and the solvent mask have opposite handedness relative to `emd_8731.mrc`. However, this does not affect *spIsoNet* training. Users may choose to flip the handedness of the half-maps and solvent mask to match `emd_8731.mrc`. In this protocol, we instead flip the final post-processed map after *spIsoNet* map Anisotropy Correction using the ChimeraX command `vop flip #vol_num`, where `vol_num` should be replaced with the actual volume number.

### ? TROUBLESHOOTING

**3DFSC computation** ● Timing ∼2 min

2 Calculate the 3DFSC volume. The default output file name is `FSC3D.mrc`, which describes the Fourier shell correlation (FSC) in different directions (Fig. 2e). This step does not require GPU acceleration. Multiple CPU cores can be used for parallelization by specifying `--ncpus`. The resulting `FSC3D.mrc` file will be used as the `aniso_file` in the subsequent *spIsoNet* reconstruction step. The inputs for 3DFSC calculation are two unfiltered half-maps and a solvent mask. Users can also set the resolution limit for 3DFSC calculation by specifying `--limit_res`; for the example dataset used here, this is set to `--limit_res 3.5`. By default, this value is set to the overall resolution of the cryoEM map on the basis of the gold-standard FSC criterion of 0.143. Run the following command:

`spisonet.py fsc3d emd_8731_half_map_1.mrc`

`emd_8731_half_map_2.mrc emd_8731_msk_1.mrc --ncpus 16`

`--limit_res 3.5`

### PAUSE POINT

Regions with low intensity values in an anisotropic 3DFSC volume (Fig. 2f) indicate under-sampled regions in Fourier space, whereas an isotropic map will generate an isotropic 3DFSC volume. Therefore, we use 3DFSC as a proxy for directional resolution and map anisotropy. Applying an anisotropic 3DFSC volume as a filter to a map in Fourier space approximates a cryoEM map with uneven orientation sampling.

### ? TROUBLESHOOTING

**Training the *spIsoNet* network** ● Timing ∼35 min, depending on the hardware configuration, the number of GPUs used, and the number of training epochs

3 Training the *spIsoNet* network for map Anisotropy Correction from scratch. A description of all available training parameters can be displayed by running `spisonet.py reconstruct --help`. The following parameters are critical and may need to be adjusted for different cryoEM map datasets.

##### Critical Parameters

`--alpha` %% the relative weighting between the data-consistency loss and the rotation-equivariance loss. The default value is 1. A larger value places greater weight on the rotation-equivariance loss and generally produces smoother maps.

`--beta` %% the relative weighting between denoising and missing-information recovery. A larger value places greater weight on denoising.

`--limit_res` %% the resolution limit for *spIsoNet* training during map Anisotropy Correction. By default, this value is set to the overall resolution of the cryoEM map as determined by the gold-standard FSC criterion.

`--epochs` %% the number of epochs used to train the neural network. The default value is 30.

Run the following command:

`spisonet.py reconstruct emd_8731_half_map_1.mrc`

`emd_8731_half_map_2.mrc --aniso_file FSC3D.mrc –mask`

`emd_8731_msk_1.mrc --limit_res 3.5 --epochs 30 –alpha 1 --beta 0.5 --output_dir isonet_maps --gpuID 0,1,2,3 --acc_batches 2`

### PAUSE POINT

This step creates a folder to store the output files from *spIsoNet* correction. The anisotropy-corrected maps are saved as `correctedXXX.mrc` files in the output folder. The trained neural network model, `XXX.pt`, and the loss curve, `lossXXX.png`, are also saved in this folder.

### ? TROUBLESHOOTING

4 Continue training from a pretrained model or perform direct prediction. If a job is interrupted, or if users want to restart or extend training, training can be resumed from a saved model by specifying `--pretrained_model`. When starting from a pretrained model, users may also adjust the number of epochs. For example, if the pretrained model was saved after 30 epochs, setting `–-epochs 20` continues training for 20 additional epochs, equivalent to 50 total epochs from scratch. Run the following command:

spisonet.py reconstruct emd_8731_half_map_1.mrc emd_8731_half_map_2.mrc --aniso_file FSC3D.mrc --mask emd_8731_msk_1.mrc --limit_res 3.5 --epochs 20 –alpha 1 --beta 0.5 --output_dir spisonet_continued_maps --gpuID 0,1,2,3 --acc_batches 2 --pretrained_model spisonet_maps/XXX.pt

5 Users can also set `–epochs` to 0 together with `--pretrained_model` to perform prediction only, without further training. Run the following command:

`spisonet.py reconstruct emd_8731_half_map_1.mrc`

`emd_8731_half_map_2.mrc --aniso_file FSC3D.mrc –mask`

`emd_8731_msk_1.mrc --limit_res 3.5 --epochs 0 –alpha 1 --beta 0.5 --output_dir`

`spisonet_directprediction_maps --gpuID 0,1,2,3 --acc_batches 2 --pretrained_model spisonet_maps/XXX.pt`

### ? TROUBLESHOOTING

**Map post-processing** ● Timing ∼2 min

6 Perform RELION post-processing. Post-processing of the anisotropy-corrected half-maps is not implemented in *spIsoNet*. In this protocol, we use `relion_postprocess` in RELION for B-factor estimation and map sharpening. Run the following command:

`relion_postprocess –i`

`corrected_emd_8731_half_map_1.mrc --i2`

`corrected_emd_8731_half_map_2.mrc –mask`

`emd_8731_msk_1.mrc --auto_bfac`

Alternatively, users can define their own B-factor and low-pass filtered values by running:

`relion_postprocess –i`

`corrected_emd_8731_half_map_1.mrc --i2`

`corrected_emd_8731_half_map_2.mrc --mask mask.mrc –-`

`adhoc_bfac <USER defined bfactor> --low_pass <user`

`defined resolution>`

The post-processed map has different handedness relative to the deposited EMDB map, `emd_8731.mrc`. Users can flip the post-processed map using the ChimeraX command `vop flip #vol_num`, where `vol_num` should be replaced with the actual volume number.

### PAUSE POINT

Noise2Noise-based denoising is used by default during the *spIsoNet* Anistropy Correction process and produces cleaner maps. Denoised *spIsoNet* output maps are intended for visualization and model interpretation. Users can inspect the loss curve, run *spIsoNet* correction with different numbers of epochs, and perform map post-processing with the same B-factor. The resulting maps can then be opened in IMOD, allowing users to rapidly load, inspect, and compare map quality across multiple outputs. This comparison can help determine the optimal result. For the example dataset used here, the 30-epoch result is satisfactory, and further training to 50 epochs does not produce substantial differences in map quality (Extended Data Fig. 1).

### ? TROUBLESHOOTING

**Procedure 2| Particle Misalignment Correction** ●Timing ∼6h

**Preparation: arrangement of particle-stack input files and directories** ●Timing ∼3 min

7 Misalignment Correction requires the installation of *spIsoNet* and RELION version 3.1 or 4.0. In this procedure, we assume that the base directory contains the downloaded subset of EMPIAR-10096 available at https://zenodo.org/records/20783378. This subset includes the particle metadata file with CTF information (`job025_tutorial.star`), the particle stack (`job025_tutorial.mrcs`), the mask file (`mask.mrc`), and a low-resolution reference map at 10 Å resolution (`HA_reference.mrc`), located in `./spIsonet_misalignment/EMPIAR-10096/`.

The majority of the commands in this procedure are run within this base directory. Ensure that the *spIsoNet* conda environment is activated, as it is required for all *spIsoNet* commands. Enter the following commands at the terminal:

`cd /path/to/base/directory`

`conda activate spisonet`

`spisonet.py --help`

### ? TROUBLESHOOTING

**Set the RELION external reconstruction environment** ●Timing ∼2 min

8 Set the `RELION_EXTERNAL_RECONSTRUCT_EXECUTABLE` environment variable to point to `relion_wrapper.py`. Enter the following commands at the terminal:

`export RELION_EXTERNAL_RECONSTRUCT_EXECUTABLE="python <PATH to spIsoNet>/spIsoNet/bin/relion_wrapper.py"`

`export CONDA_ENV="spisonet"`

`export PATH=<PATH to spIsoNet>/spIsoNet/bin:$PATH`

`export PYTHONPATH=<PATH to spIsoNet>:$PYTHONPATH`

9 Execute the `relion_wrapper.py` script during iterative RELION refinement. To call `relion_wrapper.py` during `relion_refine`, add the argument `--external_reconstruct` either to the command line or to the Additional arguments field under the Running tab in RELION GUI (Extended

Data Fig. 2). To use *spIsoNet* Misalignment Correction, the RELION refinement command should include the following arguments:

1. --`external_reconstruct`

2. --`solvent_mask`

3. --`solvent_correct_fsc`

4. --`keep_lowres (optional)`

### PAUSE POINT

In cryoEM datasets with severe preferred orientation, standard RELION refinement may fail to achieve reliable particle alignment during iterative refinement. In such cases, `--keep_lowres` is necessary to preserve low-resolution information from a user-provided reliable reference throughout subsequent RELION iterative alignment. The low-resolution cutoff to be retained is defined by the `--ini_high` parameter in RELION. The low-resolution reference map can be obtained using various approaches, such as reconstruction from a small particle dataset collected with stage tilt or subtomogram averaging from tomography datasets.

### ? TROUBLESHOOTING

**Anisotropy correction-powered particle Misalignment Correction** ●Timing ∼6 h, depending on the hardware configuration, the number of GPUs used, and the number of training epochs

10 Re-estimate particle poses in RELION using *spIsoNet*. In this protocol, RELION version 4.0 is used. Start a new RELION refinement from the baseline run directory and enable external reconstruction. *SpIsoNet* trains and applies anisotropy correction internally during each iteration of RELION refinement. This step is expected to require substantially longer runtime than standard RELION refinement. For the EMPIAR-10096 subset used in this protocol, the estimated processing time is approximately 6 h using four NVIDIA A100 GPUs. Additional Linux environment variables can be set as described below to tune the key parameters of *spIsoNet* Misalignment Correction.

##### Critical Parameters

`export CUDA_VISIBLE_DEVICES=”0,1,2,3”` %% Specifies which GPU(s) to use. By default, *spIsoNet* uses all available GPUs.

`export ISONET_ALPHA=1` %% the relative weighting between the data-consistency loss and the rotation-equivariance loss. The default value is 1. A larger value places greater weight on the rotation-equivariance loss and generally produces smoother maps.

`export ISONET_BETA=0.5` %% the relative weighting between denoising and missing-information recovery. A larger value places greater weight on denoising.

`export ISONET_START_HEALPIX=3` %% Defines the angular sampling step at which *spIsoNet* training starts.

`export ISONET_START_RESOLUTION=15` %% Defines the refinement resolution at which *spIsoNet* training starts.

`export ISONET_RETRAIN_EACH_ITER=True` %% Defines whether the network is retrained from scratch at each RELION refinement iteration. This is typically set to True.

`export ISONET_EPOCHS=5` %% Defines the number of epochs used to train the neural network.

`export ISONET_KEEP_LOWRES=False` %% Defines whether low-resolution information from a correct reference map is retained during particle alignment. This parameter can be overridden by the `--keep_lowres` parameter in the RELION command line or GUI.

`export ISONET_WHITENING=True` %% Defines whether whitening is performed before running *spIsoNet*.

`export ISONET_WHITENING_LOW=10` %% Defines the starting resolution for whitening.

`export ISONET_FSC_WEIGHTING=True` %%Defines whether FSC weighting is performed before running *spIsoNet*. This is typically set True.

Run the following command:

`mpirun -np 5 ‘which relion_refine_mp’ --o`

`Refine3D/job001/run --auto_refine –`

`split_random_halves --i job025_tutorial.star –ref`

`HA_reference.mrc --firstiter_cc --ini_high 10 --`

`dont_combine_weights_via_disc --preread_images --pool`

`30 --pad 2 --ctf --particle_diameter 170 --flatten_solvent --zero_mask --solvent_mask mask.mrc --solvent_correct_fsc --oversampling 1 --healpix_order 2`

`--auto_local_healpix_order 5 --offset_range 5 --offset_step 2 --sym C3 --low_resol_join_halves 40 --norm --scale --j 4 --gpu "" --external_reconstruct --keep_lowres --pipeline_control Refine3D/job001/`

### PAUSE POINT

For this particle dataset, which exhibits severe preferred orientation, we retained low-resolution information to 10Å from the reference by setting --keep_lowres, thereby improving the reliability of particle alignment.

### ? TROUBLESHOOTING

2 Misalignment correction generates intermediate anisotropy-corrected half-maps at each iteration of RELION refinement for particle pose estimation, These maps are saved as correctedXXX.mrc files in the refinement output directory.

Users can also inspect the RELION-generated half-maps from each iteration (`run_itXXX_half1_unfil_backup.mrc`), which are reconstructed based on the updated particle Euler angles, to evaluate the effect of misalignment correction (Extended Data Fig. 3). After the refinement is complete, RELION generates the final half-maps (`run_half1_class001_unfil.mrc)` and (`run_half2_class001_unfil.mrc)` and the refined particle metadata file (`run_data.star`).

### PAUSE POINT

The 3DFSC volume, the trained neural network model (XXX.pt), and a plot showing the training loss (lossXXX.png) are also generated for each iteration.

### ? TROUBLESHOOTING

#### Map post-processing

3 Perform post-processing of the final half-maps generated by RELION. In this procedure, `relion_postprocess` is used for map sharpening. Enter the following command:

`relion_postprocess --i run_half1_class001_unfil.mrc --i2 run_half2_class001_unfil.mrc --mask mask.mrc --auto_bfac`

Alternatively, users can define their own B-factor and low-pass filtered resolution by running:

`relion_postprocess --i run_half1_class001_unfil.mrc --i2 run_half2_class001_unfil.mrc --mask mask.mrc --`

`adhoc_bfac <USER defined bfactor> --low_pass <USER defined resolution>`

### PAUSE POINT

Because the *spIsoNet* misalignment-corrected half-maps are independently reconstructed in RELION using the refined particle poses in run_data.star, their independence is preserved, allowing them to be used for gold-standard FSC-based overall resolution estimation.

### CRITICAL

Users can optionally use these misalignment-corrected half-maps for further map Anisotropy Correction following Procedure 1, thereby addressing both major consequences of preferred orientation.

### Troubleshooting

User questions and feedback were collected through the *spIsoNet* Google Group: https://groups.google.com/u/0/g/spisonet. Troubleshooting information is provided in Table 2.

**Table 2.**
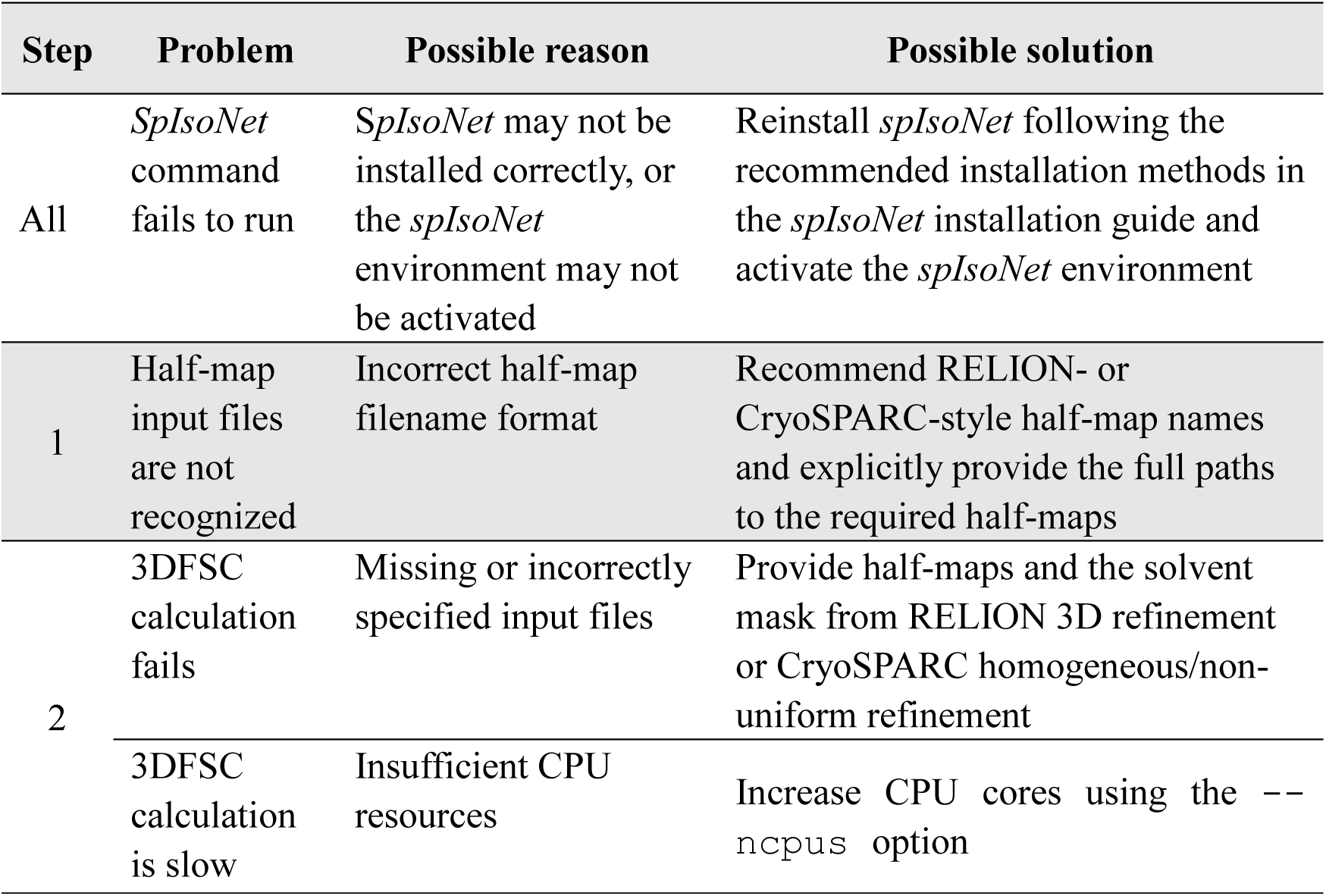
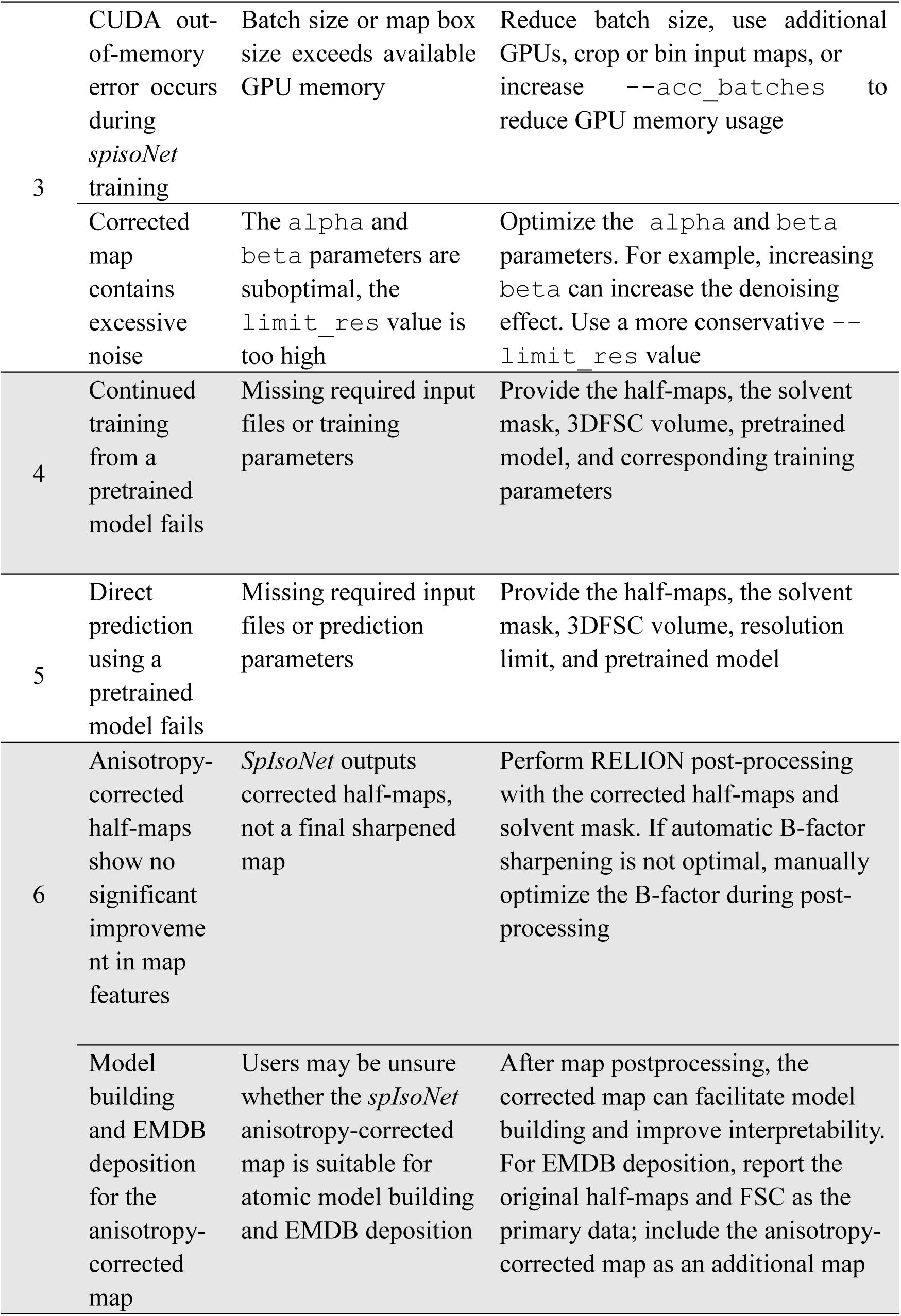
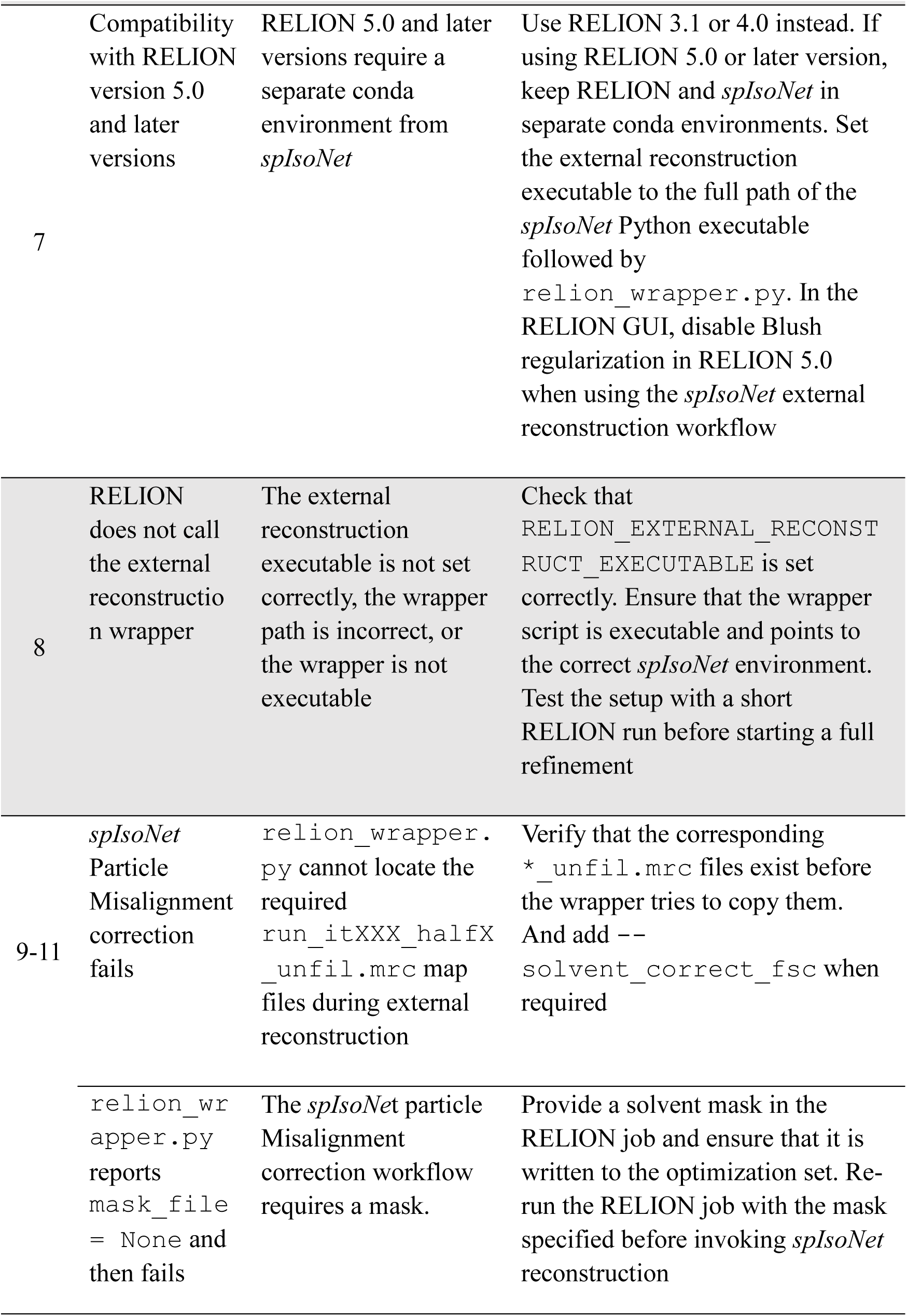
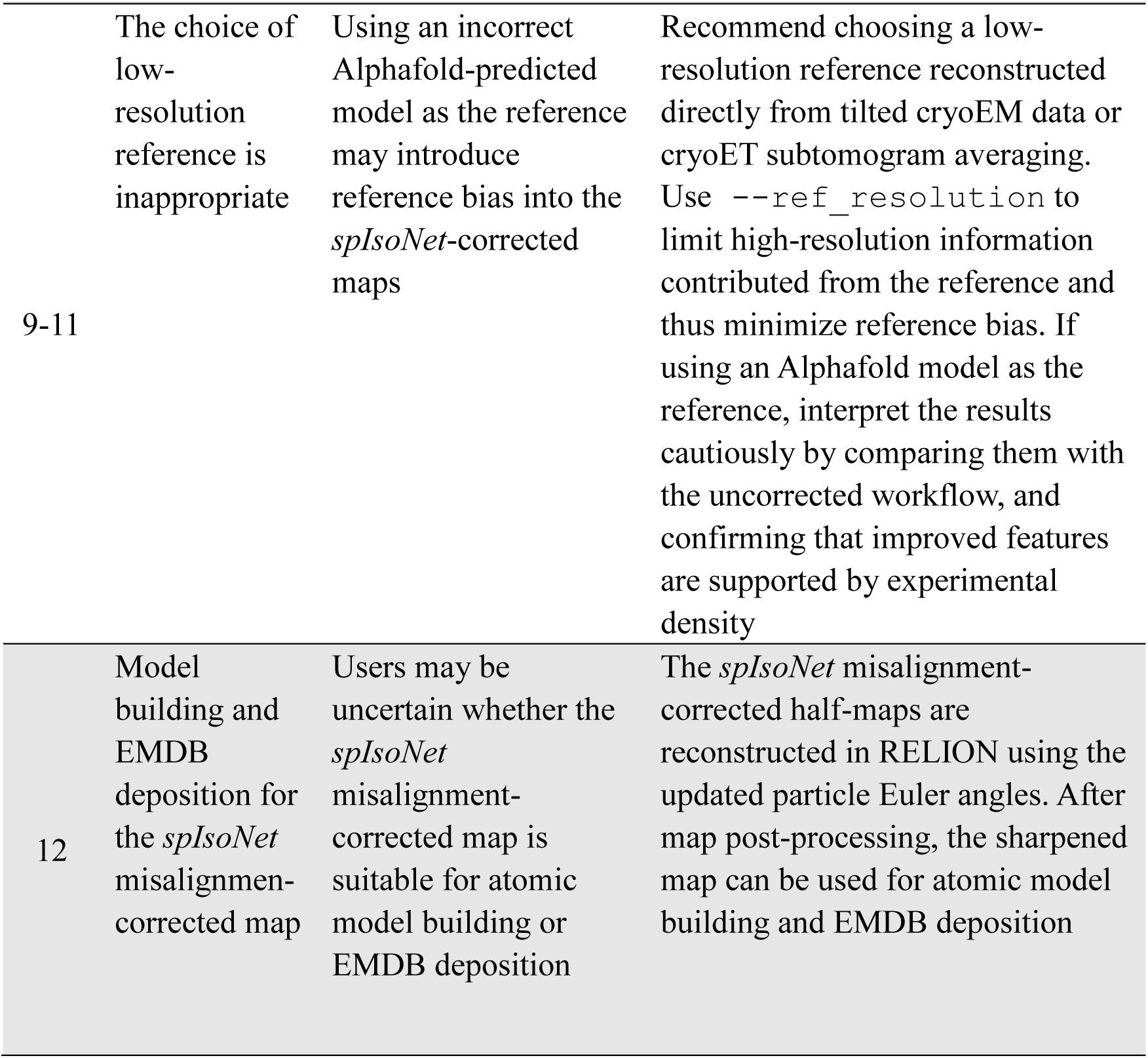
Troubleshooting table.

### Timing

The time required to run this protocol depends on the available hardware. We provide approximate timings in each section of the protocol based on our hardware described in Materials. For users applying this protocol to their own datasets, runtime is mainly determined by the duration of *spIsoNet* model training and the size of the particle images, as these steps are the most demanding in terms of both time and computational resources. Instructions for how to use *spIsoNet* are also available at https://github.com/IsoNet-cryoET/spIsoNet/tree/main/tutorial.

### Anticipated results

We illustrate the anticipated results of this protocol using two test datasets: (i) deposited half-maps reconstructed from a tilted cryoEM influenza hemagglutinin (HA) trimer dataset for Procedure 1 and (ii) an untilted cryoEM influenza HA trimer particle dataset for Procedure 2.

In the first test dataset, the deposited cryoEM map, EMD-8731, reconstructed using standard RELION Refinement, shows relatively poor map quality due to preferred orientation, as reflected by rod-like α-helical densities with weak or missing side-chain densities (Fig. 3a-b). We downloaded the EMD-8731 half-maps and solvent mask and applied *spIsoNet* map Anisotropic Correction to the half-maps, followed by map post-processing. Compared with the deposited `emd_8731.mrc` map, the anisotropy-corrected map exhibits a more clearly resolved α-helical pitch and improved side-chain densities (Fig. 3c-d).

**Fig. 3.**
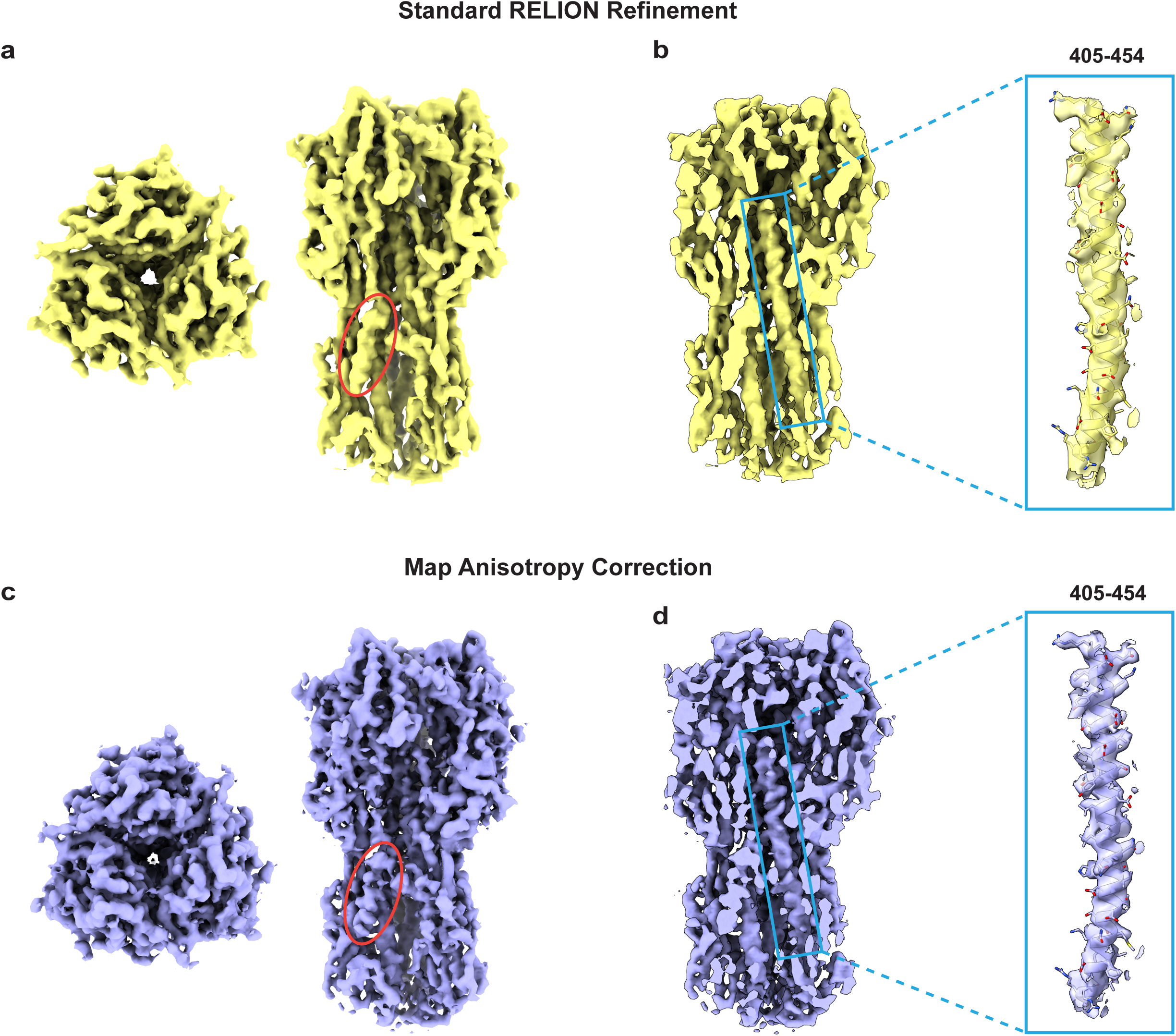
*SpIsoNet* map Anisotropy Correction of the cryoEM map reconstructed from the tilted influenza HA trimer dataset (EMD-8731). **a**, CryoEM map of the HA trimer obtained by standard RELION Refinement. **b**, Representative α-helical density of the HA trimer fitted with the fitted atomic model. Residue numbers are indicated. **c**, CryoEM map of the HA trimer after *spIsoNet* map Anisotropy Correction. **d**, Representative α-helical density of the HA trimer with the atomic model fitted after *spIsoNet* correction. Residue numbers are indicated.

In the second test dataset, a representative cryoEM micrograph and RELION 2D classification results show that the particle dataset consists predominantly of top views, with rare side views observed in the 2D class averages (Fig. 4a-b). This HA trimer particle dataset therefore exhibits severe preferred orientation, resulting in a highly distorted cryoEM map with no interpretable secondary-structure features when processed using the standard RELION Refinement pipelines (Fig. 4c). We applied *spIsoNet* Misalignment Correction during RELION 3D auto-refinement. Particle orientations were first re-estimated through a global orientation search followed by a local orientation search and map post-processing, yielding an approximately 3.5 Å reconstruction (Fig. 4d–f, Extended Data Fig. 3). Typical structural features, including α-helices, β-sheets, and side-chain densities, are clearly visible in this map.

**Fig. 4.**
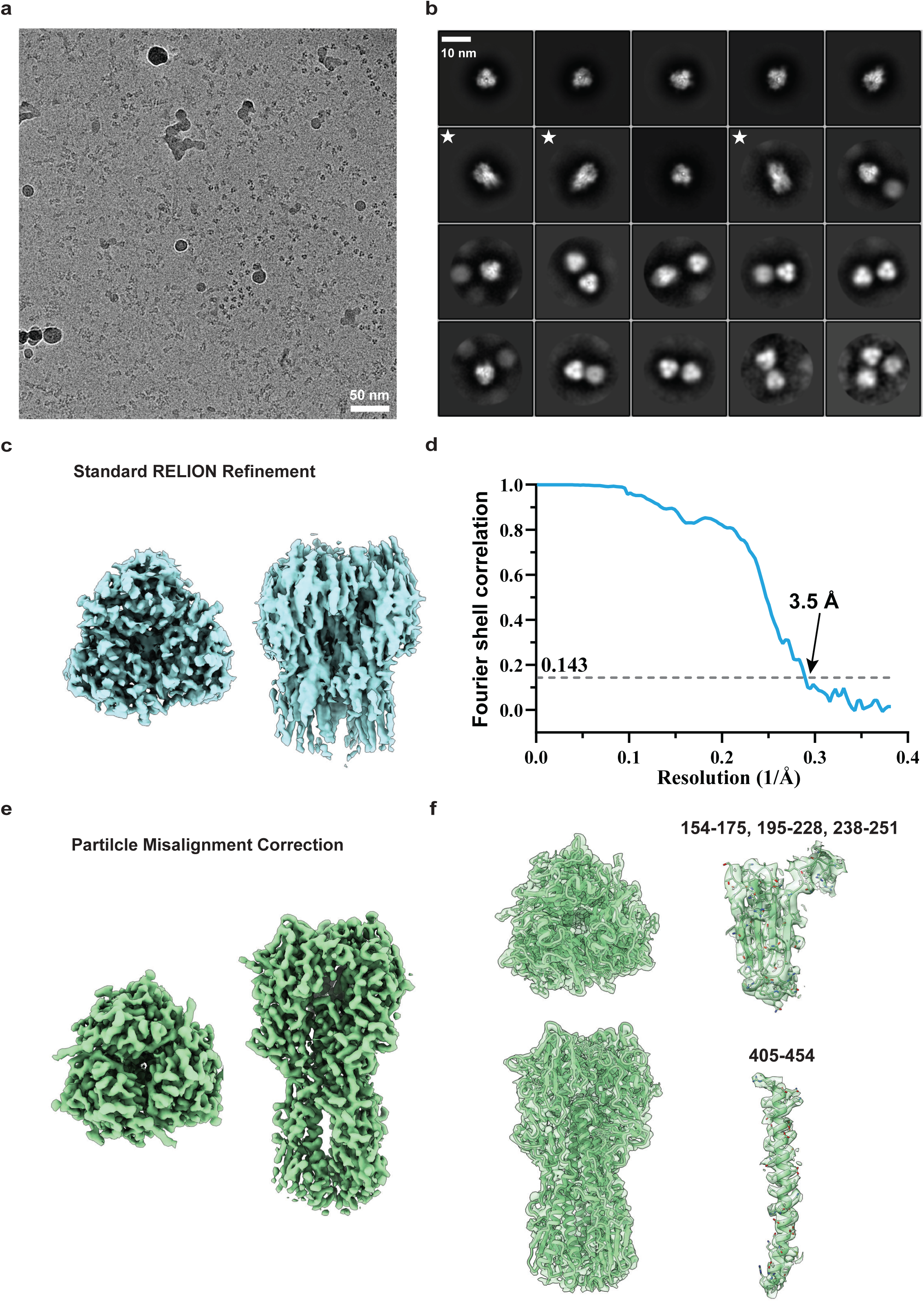
*SpIsoNet* particle Misalignment Correction of the untilted HA trimer particle images (EMPIAR-10096). **a**, Representative cryoEM micrograph of the untilted influenza HA trimer dataset. **b**, Representative 2D class averages computed from the EMPIAR-10096 particle subset. Oblique and side views are indicated by white stars. **c**, CryoEM map of the HA trimer obtained by standard RELION Refinement. **d**, Gold-standard FSC curve of the HA trimer map after Misalignment Correction. **e**, CryoEM map of the HA trimer after *spIsoNet* particle Misalignment Correction. **f**, Model fitting and representative density regions of the HA trimer after *spIsoNet* particle Misalignment Correction, shown with the fitted atomic model. Residue numbers are indicated.

Overall, by following this protocol, users should be able to reproducibly generate the expected outputs, including:

1. trained *spIsoNet* models for prediction;
2. a *spIsoNet* training-loss curve plot reflecting model training performance;
3. *spIsoNet* anisotropy-corrected half-maps generated using Procedure 1.
4. *spIsoNet* particle misalignment-corrected half-maps generated through RELION refinement using Procedure 2.

## Reporting summary

Further information on research design is available in the Nature Research Reporting Summary linked to this article.

## Data availability

All expected results presented in this protocol are available at https://zenodo.org/records/20783378.

## Code availability

The *spIsoNet* is available at the GitHub under MIT license (https://github.com/IsoNet-cryoET/spIsoNet).

## Acknowledgements

This project is supported by a grant from the US National Institutes of Health (R01GM071940 to Z.H.Z.) and the American Heart Association (AHA) Postdoctoral Fellowship (26POST1570844 to H.F.).

## Author contributions

H.F. and Y.-T.L. contributed to preparing the *spIsoNet* workflow protocol, writing the paper and preparing the figures. Z.H.Z. oversaw the project and edited the manuscript.

## Additional information

The full version comes with one supplementary file: the *spIsoNet* tutorial document.

---

**Extended Data Fig. 1.**
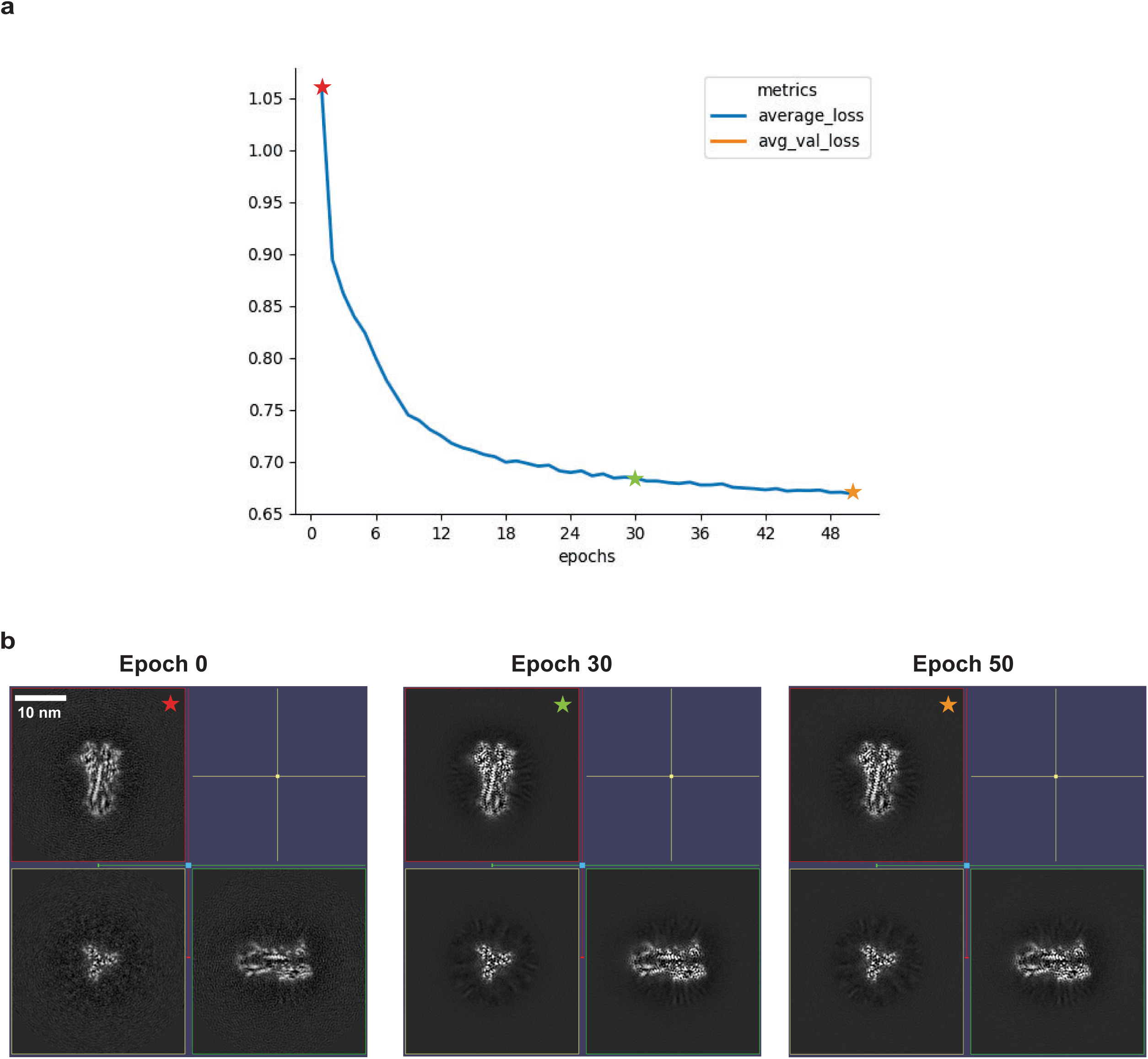
Training process for *spIsoNet* map Anisotropy Correction. **a**, Loss curve during *spIsoNet* training. **b**, Orthogonal slice through cryoEM maps from epoch 0, 30 and 50. All map were sharpened using the same parameters.

**Extended Data Fig. 2.**
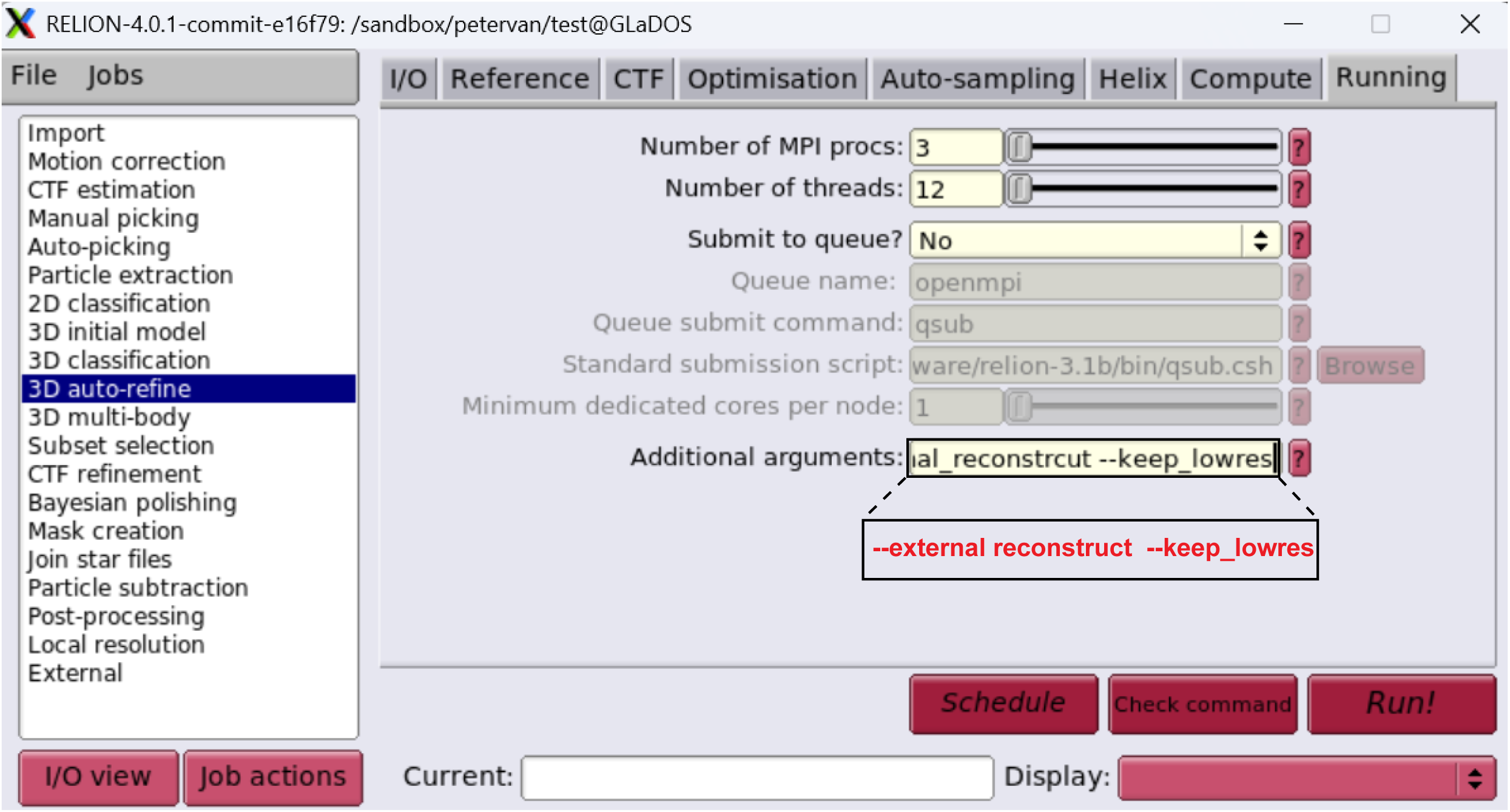
Configuration of *spIsoNet* particle Misalignment Correction in the RELION 4 graphical user interface (GUI). Add the `--external_reconstruct` parameter and, optionally, the `--keep_lowres` parameter.

**Extended Data Fig. 3.**
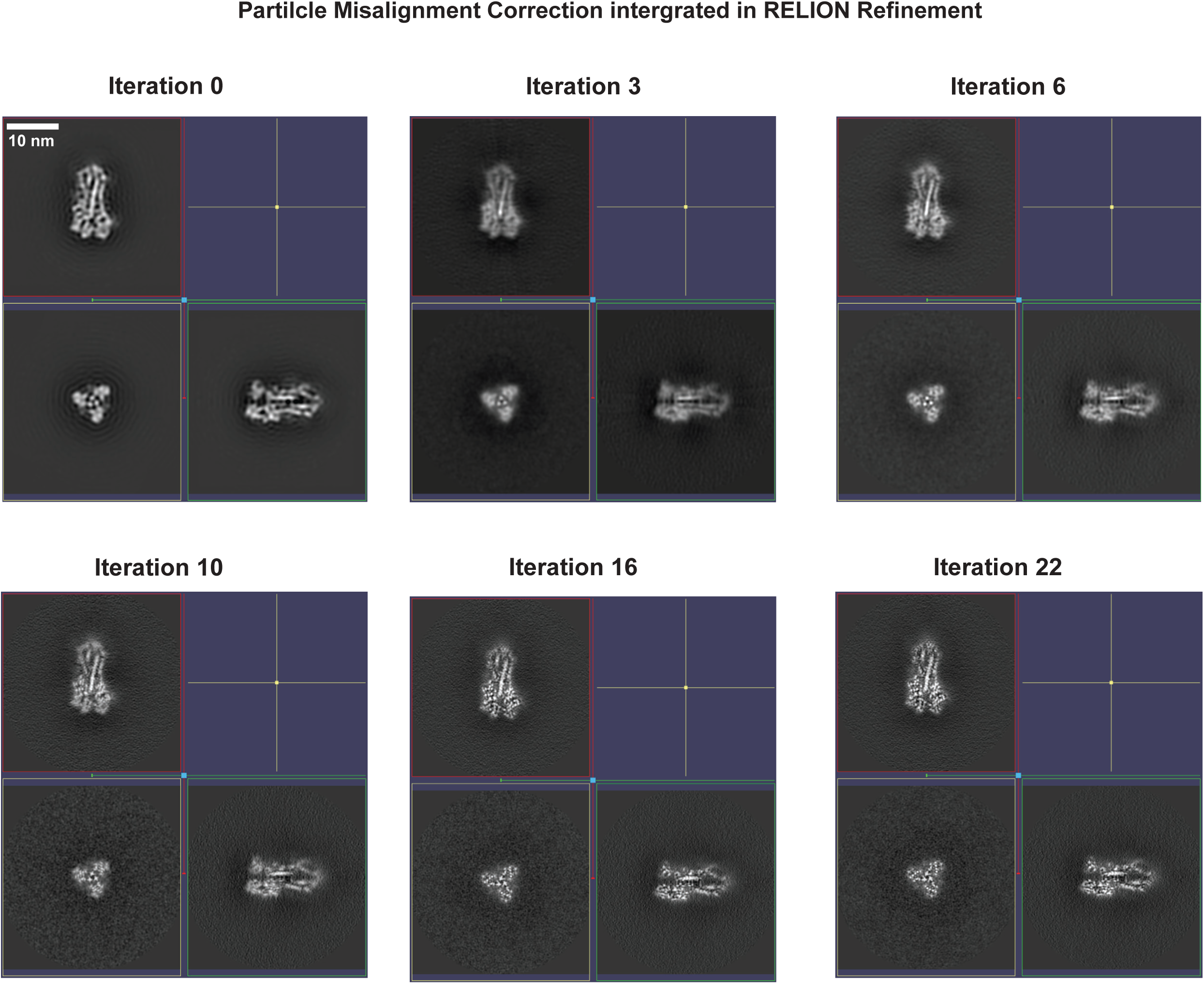
Iterative refinement process for *spIsoNet* particle Misalignment Correction. Orthogonal slices through cryoEM half-maps reconstructed in RELION at iterations 0, 3, 6, 10, 16, and 22.

